# Membrane receptor MerTK is a newly identified transcriptional regulator that associates to chromatin as nanoclusters during human DC differentiation

**DOI:** 10.1101/2020.04.16.044974

**Authors:** Kyra. J.E. Borgman, Georgina Flórez-Grau, Maria A. Ricci, Carlo Manzo, Melike Lakadamyali, Alessandra Cambi, Daniel Benítez-Ribas, Felix Campelo, Maria. F. Garcia-Parajo

## Abstract

MerTK is a transmembrane receptor tyrosine kinase (RTK) mainly expressed in dendritic cells (DCs) and macrophages where it plays an important role in immunotolerance, but also in activating oncogenic signalling pathways. Albeit MerTK is exploited as clinical target in cancer and auto-immune disorders, the mechanisms that regulate its diverse functions are poorly understood. Here, we identified a remarkably high pool of the full receptor in the nucleus of human DCs. Nuclear translocation was ligand-dependent. Importantly, MerTK nuclear levels correlated to DC differentiation and were spatiotemporally regulated by the transmembrane receptor LRP-1. Using dual-colour super-resolution nanoscopy we discovered that nuclear MerTK forms nanoclusters, whose strength strongly depends on chromatin accessibility during DC differentiation. We finally revealed high transcription capacity of MerTK. Overall, our work indicates that nuclear MerTK acts as a transcription factor regulating DC differentiation, thus implicating for the first time a physiological function for RTK nuclear translocation in immunity.

## Introduction

RTKs comprise a family of cell-surface receptors key in regulating essential cellular processes such as growth, differentiation, survival and migration (Lemmon and Schlessinger, 2010). Structurally highly conserved, these receptors contain an extracellular domain, a single transmembrane domain, and an intracellular kinase domain. Ligand binding at the extracellular domain activates the receptor by inducing homo-dimerization and subsequent auto-phosphorylation of tyrosine residues in the cytoplasmic tail that initiate downstream signalling cascades (Hubbard, 1999; Li and Hristova, n.d.). MerTK is a member of the RTK family that regulates an intriguingly broad range of seemingly unrelated cellular processes, including apoptosis, migration, transcription (Cummings et al., 2013; Graham et al., 2014; Linger et al., 2008), and immunotolerance (Cabezón et al., 2015; Camenisch et al., 1999; Lu and Lemke, 2001; Rothlin et al., 2007; Rothlin and Lemke, 2010; Scott et al., 2001; Wallet et al., 2008). Physiologically, MerTK is mainly expressed on the surface of macrophages and DCs (Behrens et al., 2003), where it plays a role in phagocytosis of apoptotic cells (Scott et al., 2001) and in downregulating the secretion of pro-inflammatory cytokines (Sen et al., 2007). Loss of MerTK function and of its family members Tyro-3 and Axl (TAM family) leads to inflammation and increased susceptibility for auto-immune disorders (Lu and Lemke, 2001; Rothlin and Lemke, 2010). In contrast, ectopic or increased expression of MerTK has been found in a wide variety of cancers where it activates oncogenic signalling pathways leading to increased cell survival, invasion, and therapy resistance (Cummings et al., 2013; Graham et al., 2014).

Due to its broad involvement in cancer and auto-immune disorders, MerTK is being increasingly exploited as a potential clinical target. Multiple reports have demonstrated the effectiveness and specificity of MerTK inhibition in tumour suppression (Brandao et al., 2013; Cook et al., 2013; Crittenden et al., 2016; Cummings et al., 2015; Kim et al., 2017). In the context of immunity, a recent study in human DCs showed that MerTK is highly upregulated upon several days of tolerogenic treatment with Dexamethasone (Cabezón et al., 2015). These so-called tolerogenic DCs suppress both T cell expansion and pro-inflammatory cytokine production by T cells (Cabezón et al., 2015), a process that is regulated by MerTK. Several ongoing clinical trials indeed exploit the immune tempering function of MerTK, among other immunosuppressive mechanisms, by injecting tolerogenic DCs into patients in order to battle different auto-immune disorders such as diabetes type I (Giannoukakis, 2013), Rheumatoid Arthritis (Bell et al., 2017; Benham et al., 2015) or Crohn’s disease (Jauregui-Amezaga et al., 2015). Surprisingly, despite the evident clinical relevance of MerTK signalling, very little is known on the molecular mechanisms of action by which this receptor is able to accomplish its broad range of functions.

Although the function of MerTK has been classically associated to its expression at the plasma membrane, a recent study on human tolerogenic DCs identified an abnormally large pool of the receptor located intracellularly and accounting for as much as 40% of its total expression levels (Cabezón et al., 2015). However, the subcellular location as well as function of this intracellular MerTK pool has remained completely elusive. We hypothesized that the existence of several pools of the receptor with distinct subcellular localizations might be important in defining its functional diversity. We thus employed biochemical tools together with advanced optical imaging techniques, including super-resolution microscopy, to investigate the spatial distribution of MerTK in immunogenic and tolerogenic human DCs. Remarkably, we found that intracellular MerTK is mainly located in the nucleus and that its degree of nuclear accumulation is strictly related to DC differentiation. Moreover, once in the nucleus, MerTK associates to chromatin and it is capable to induce transcription. As a whole, our results indicate that aside from its well-established role on the cell membrane, the residence of MerTK in the nucleus constitutes a physiological relevant mechanism for dendritic cells, functioning as a potential genomic regulator during DC differentiation. Given the involvement of MerTK in both auto-immunity and cancer, our results might have impact on the broad implementation of MerTK for clinical therapy purposes.

## Results

### MerTK is found both on the membrane and in the nucleus of tolerogenic DCs

Previous studies by Cabezon et al. (Cabezón et al., 2015) showed that MerTK is highly upregulated in immature tolerogenic DCs upon several days of tolerogenic treatment with the glucocorticoid Dexamethasone (Dex). We first confirmed by flow cytometry that these Dex-treated immature DCs, referred to as iDex, highly express MerTK on their membrane, as opposed to immunogenic immature DCs (iDCs) (Fig. 1A). To elucidate the spatial organization of the receptor on the cell membrane at the single cell level, we performed super-resolution, stimulated emission depletion (STED) nanoscopy imaging. With a spatial resolution of ~100nm, we identified well-separated fluorescent spots of MerTK homogeneously distributed across the plasma membrane of iDex cells (Fig. 1B). These spots correspond to small MerTK nanoclusters containing on average 3 and up to 10 receptors (see materials and methods) (Fig. 1C), and having sizes ~120nm (Fig. 1D). This kind of organization is in good agreement with the general consensus that nanoclusters are the functional unit for many immunoreceptors on the plasma membrane (Akiyama et al., 2015; Garcia-Parajo et al., 2014; Torreno-Pina et al., 2016, 2014; van Zanten et al., 2009).

**Figure 1:**
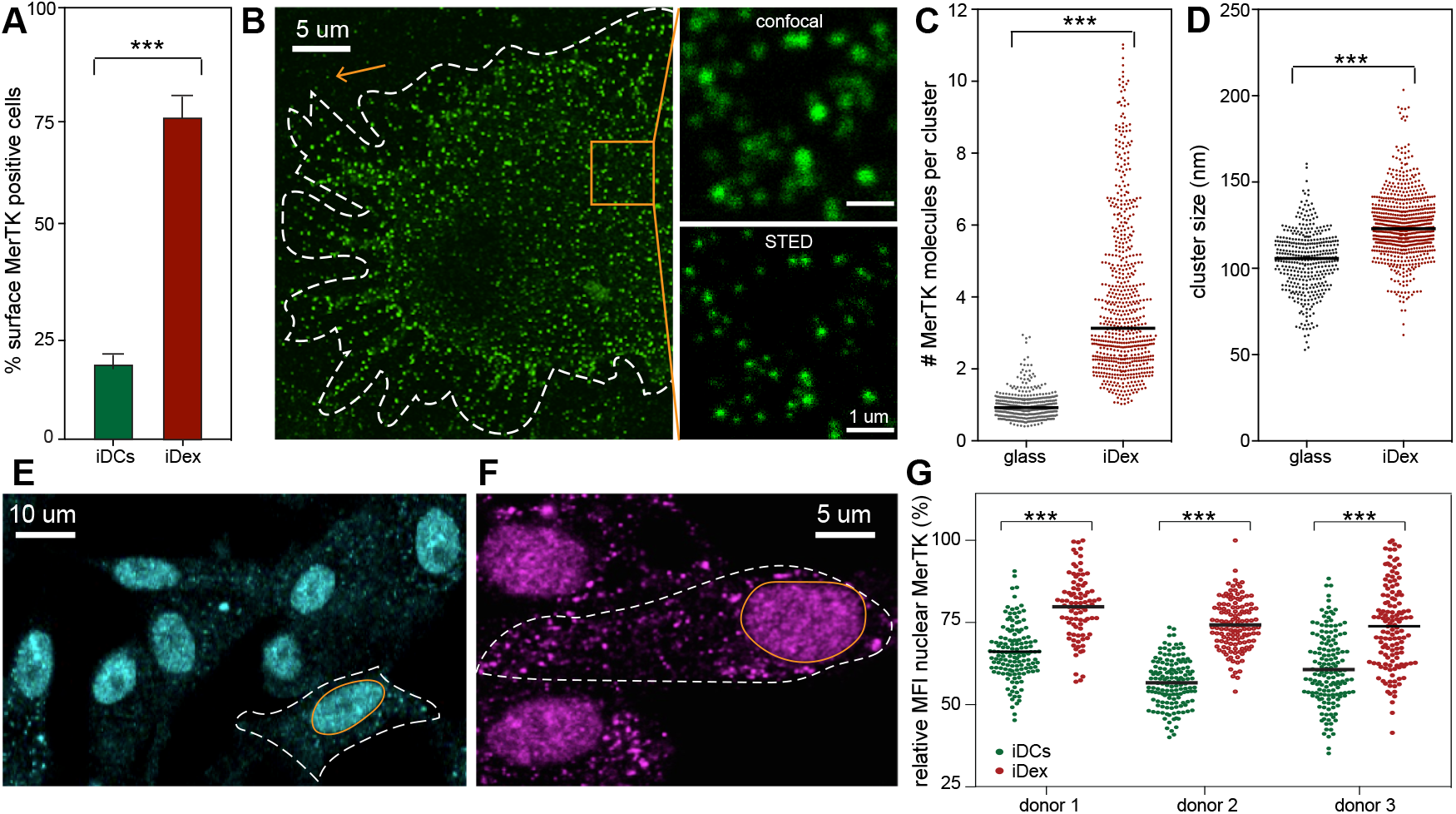
Membrane and intracellular distribution of MerTK in DCs. (**A**) Flow cytometry analysis of surface MerTK expression on iDC and iDex cells (n=8). (**B**) Representative STED image of MerTK on the plasma membrane of an iDex cell. The dotted line delineates the cell boundary. The orange square indicates the location of the zoom-in images, shown in confocal and STED modes. The orange arrow points to an individual ‘labelling unit’ on the glass that is used for the quantification in C. (**C**) Quantification of the number of MerTK molecules per nanocluster in iDexs, compared to the intensity of individual labelled antibodies on glass. (**D**) MerTK nanocluster sizes. Data from 3 different donors (around 8 cells each) for C and D. (**E**) Representative confocal image of the intracellular distribution of MerTK, using a MerTK Ab against an extracellular epitope. The dotted line represents the cell boundary, while the orange line represents the nuclear envelope. (**F**) Like in E, but using a MerTK Ab against an intracellular epitope. (**G**) Relative mean fluorescence intensity (MFI) of MerTK in the nucleus of iDCs and iDexs. Each dot corresponds to a single nucleus: Data from 80-100 cells per donor per condition.

Recent flow cytometry studies performed on human DCs showed that ~ 40% of MerTK resides intracellularly, both under immunogenic as well as tolerogenic conditions (Cabezón et al., 2015). To identify the location of this intracellular pool we performed confocal imaging of MerTK together with different organelle markers on both iDCs and iDex cells. To first exclude the possibility that the intracellular pool of MerTK corresponds to receptors targeted for degradation, we labelled MerTK and lysosomes. A clear exclusion of MerTK from the lysosome compartment was observed (Supplementary Fig. 1). Interestingly, we found that the MerTK intracellular pool almost entirely resides inside the nucleus (Fig. 1E). To validate the specificity of the antibodies used for imaging, we further probed MerTK using two other antibodies raised against different extracellular epitopes from different manufacturers, and obtained the same nuclear distribution (Supplementary Figs. 2A,B). A fourth antibody against an intracellular epitope of MerTK also gave the same spatial distribution (Fig. 1F), indicating that both intracellular and extracellular parts of the protein exhibit nuclear localization. This result also suggests that the nuclear MerTK pool observed by us, does not correspond to a previously reported soluble isoform consisting of the extracellular domain of the protein (Sather et al., 2007), nor to its intracellular counterpart that occurs after proteolytic cleavage. Quantification of the amount of nuclear MerTK in both iDCs and iDex (3 different donors, each) from fluorescent images shows that iDexs exhibit a more pronounced MerTK nuclear accumulation as compared to iDCs (Fig. 1G). Nevertheless, this increase (~20%) was much more modest as compared to the three-fold increase in the expression level detected at the cell membrane (Fig. 1A).

To further validate the physiological relevance of our results and to rule out potential artefacts caused by the *in-vitro* differentiation of the tolerogenic DCs, we isolated immune cells with a tolerogenic phenotype directly from the tumour environment. Also, in these cells, MerTK nuclear distribution was clearly observed (Supplementary Fig. 2C). Nuclear localization was detected in the monocytic cell line THP-1 as well (Supplementary Fig. 2D). Altogether, these results show a remarkable high MerTK nuclear localization in different immune cells: cells from the THP-1 cell line, *in-vitro* monocytic derived DCs, and directly isolated immune cells.

### Nuclear MerTK levels strictly correlate with DC differentiation

Membrane expression of MerTK on human DCs was previously shown to depend on tolerogenic treatment with Dex (Cabezón et al., 2015). Indeed, surface MerTK is absent in monocytes and only steeply increases in the first two days after tolerogenic treatment, in contrast to cells equally differentiated in the absence of Dex where membrane expression is always minimal (Cabezón et al., 2015). We therefore enquired whether the same dependence would hold for nuclear MerTK (*n*MerTK) upon Dex treatment. For this, we permeabilized the cells and quantified, by confocal microscopy, the amount of *n*MerTK during each of the seven days of differentiation from monocytes into iDCs/iDexs. In contrast to the strong effect that Dex had on the expression levels of MerTK on the cell membrane, tolerogenic treatment had only a minor influence on the amount of *n*MerTK, as nuclear expression levels were comparable in the presence or absence of Dex (Fig. 2A) with only a modest increase in iDex at day four (last time point in Fig. 2A, and Fig. 1G). Remarkably, a strong correlation between the amount of *n*MerTK and the stage of DC differentiation was observed, reaching maximum levels at day 0, the moment when monocytes transition into early DCs. Moreover, *n*MerTK localization occurred gradually, starting with a low overall expression in monocytes (day −2), massive expression increase and distribution throughout the entire cytoplasm (day - 1) and specific nuclear accumulation around day 0 (Fig. 2B). Nuclear accumulation then persisted as a function of DC differentiation and led to a well-defined distribution (day 4) where almost all the intracellular MerTK resides in the nucleus (Fig. 2B).

**Figure 2:**
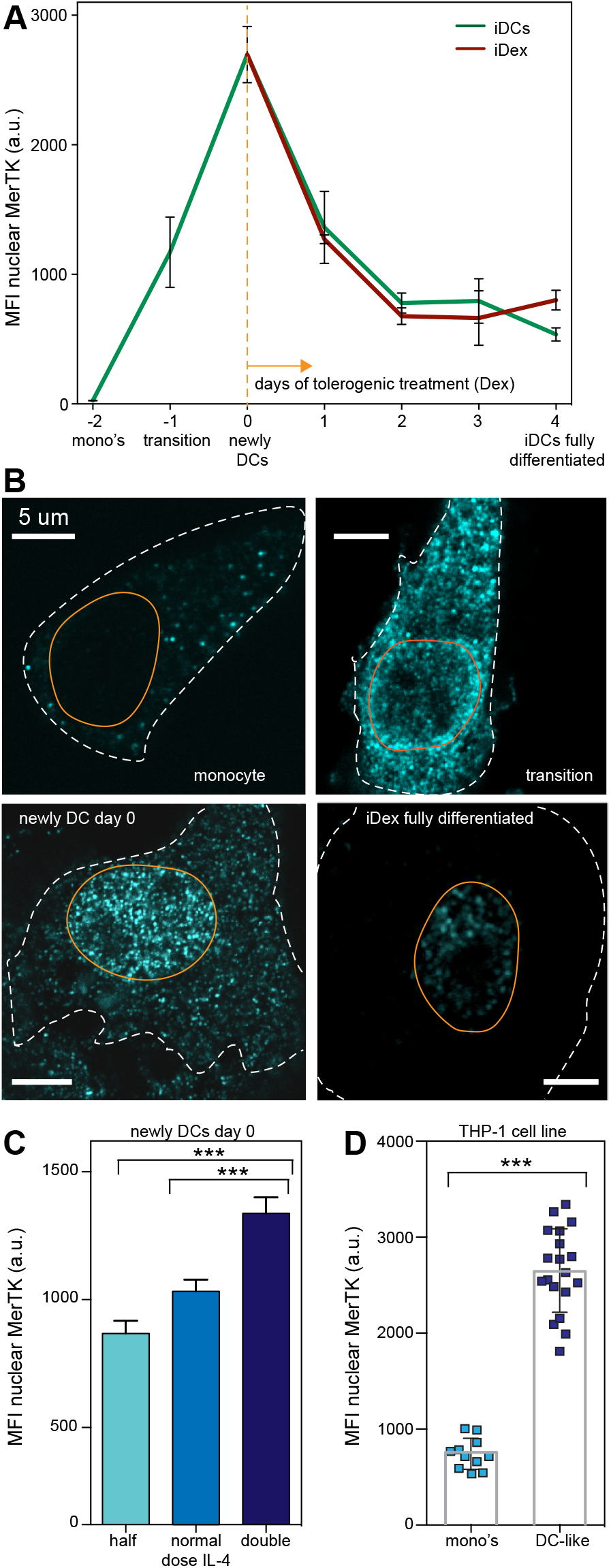
Nuclear expression of MerTK during monocyte differentiation into DCs. **(A)** MFI of *n*MerTK at different time points during monocyte differentiation into DCs. Day −2 (mono’s) corresponds to harvested monocytes, 3 hours after removal of all other leukocytes. Day −1 (transition) corresponds to 24 hours after inducing differentiation. Day 0 (newly DCs) corresponds to 48 hours after inducing differentiation. At day 0, one pool of cells were subjected to Dex treatment (iDex) and the other pool left without the treatment. For both pools, *n*MerTK was measured until fully differentiated DCs were obtained (day 4). 25-50 cells from 3 different donors per condition. (**B**) Representative confocal images of intracellular MerTK distribution in differentiating monocytes at different time points. Of note, the loss of MerTK signal at the plasma membrane is mostly due to the robust permeabilization treatment required to penetrate the nucleus. (**C**) MFI of *n*MerTK as a function of IL-4 dose during the first 2 days of differentiation (day −2 to day 0). Around 80 cells from 2 different donors per condition. (**D**) MFI of *n*MerTK in THP-1 cells before and after differentiation towards a DC-like phenotype. N=20.

To further demonstrate that this increase in *n*MerTK expression is specific to DC differentiation rather than to days of *in-vitro* culturing, we assessed the effect of the differentiation cocktail (cytokines IL-4 and GM-CSF that together facilitate the differentiation of monocytes into DCs (de Vries et al., 2005)) on the amount of *n*MerTK. First, we used different doses of IL-4 to transition monocytes into newly DCs (at day 0). As expected, the higher the dose of this differentiating cytokine (within the physiological relevant range of DC differentiation), the more MerTK accumulates into the nucleus (Fig. 2C), indicating a correlation between differentiation and *n*MerTK. Second, we administered the full cocktail to THP-1 monocytic cells that had been previously cultured for several cell cycles to induce their differentiation in to DC-like cells (Berges et al., 2005; Guo et al., 2012). Also in these conditions a strong increase in *n*MerTK was observed (Fig. 2D), along with a clear change in cell morphology indicative of a DC-like phenotype (Supplementary Fig. 3A-C). Altogether these data demonstrate that the degree of *n*MerTK expression strictly correlates with DC differentiation, reaching its maximum at the transition point from monocytes into early DCs.

Over the last decade, a few reports have indicated the presence of some RTKs in the nucleus as a malignant side effect resulting from its overexpression in tumour cells (Brand et al., 2012; Wang and Hung, 2012; Wells and Marti, 2002a). To investigate whether MerTK would follow a similar pattern, we overexpressed the receptor in different tumour cell lines. HeLa cells transfected with MerTK showed a membrane expression profile similar to that of DCs with the presence of small nanoclusters (Supplementary Fig. 3D). Nevertheless, this aberrant expression did not result in MerTK nuclear translocation (Supplementary Fig. 3E). Likewise, HEK-293 cells which endogenously express MerTK, do not exhibit *n*MerTK (Supplementary Fig. 3F), neither does its overexpression induce nuclear translocation (Supplementary Fig. 3G). Thus, in contrast to other RTKs, our data show that *n*MerTK is not the result of aberrant or overexpression of the receptor, but is rather a trait exclusively reserved for immune cells (THP-1, several types of DCs and Jurkat T cells (Migdall-Wilson et al., 2012)), suggesting a physiological role for *n*MerTK in immune cells.

### Binding of ligand ProS induces *n*MerTK translocation

Trafficking from the membrane into the nucleus has previously been reported for several RTKs (Chen and Hung, 2015; Wells and Marti, 2002a), amongst which the epidermal growth factor receptor (EGFR) has received most attention (Wang et al., 2010; Wang and Hung, 2012). In these cases, nuclear translocation was found to be ligand induced (Carpenter and Liao, 2009; Lin et al., 2001). This prompted us to explore the role of the MerTK ligand ProS (Lemke and Rothlin, 2008; Stitt et al., 1995) in *n*MerTK translocation by directly imaging the receptor and ProS on individual iDex cells. Even though GAS6 is also a well-described ligand for MerTK (Chen et al., 1997; Nagata et al., 1996), we focused on ProS as it has been previously described to be the main ligand involved in immunoregulation by human DCs (Carrera Silva et al., 2013). A strong colocalization between MerTK and ProS was observed intracellularly (Fig. 3A) together with the presence of multiple receptor-ligand complexes associated to the nuclear envelope (NE) (orange arrows in Fig. 3B). To quantify the degree of colocalization between MerTK and ProS at different subcellular regions, we segmented the cell images into periphery (mostly membrane), cytoplasm, and NE bound. At the cell periphery, colocalization is low suggesting that MerTK internalization quickly follows after ProS binding (Fig. 3C). In strong contrast, colocalization markedly increases towards the nucleus, consistent with ligand induced intracellular MerTK trafficking.

**Figure 3:**
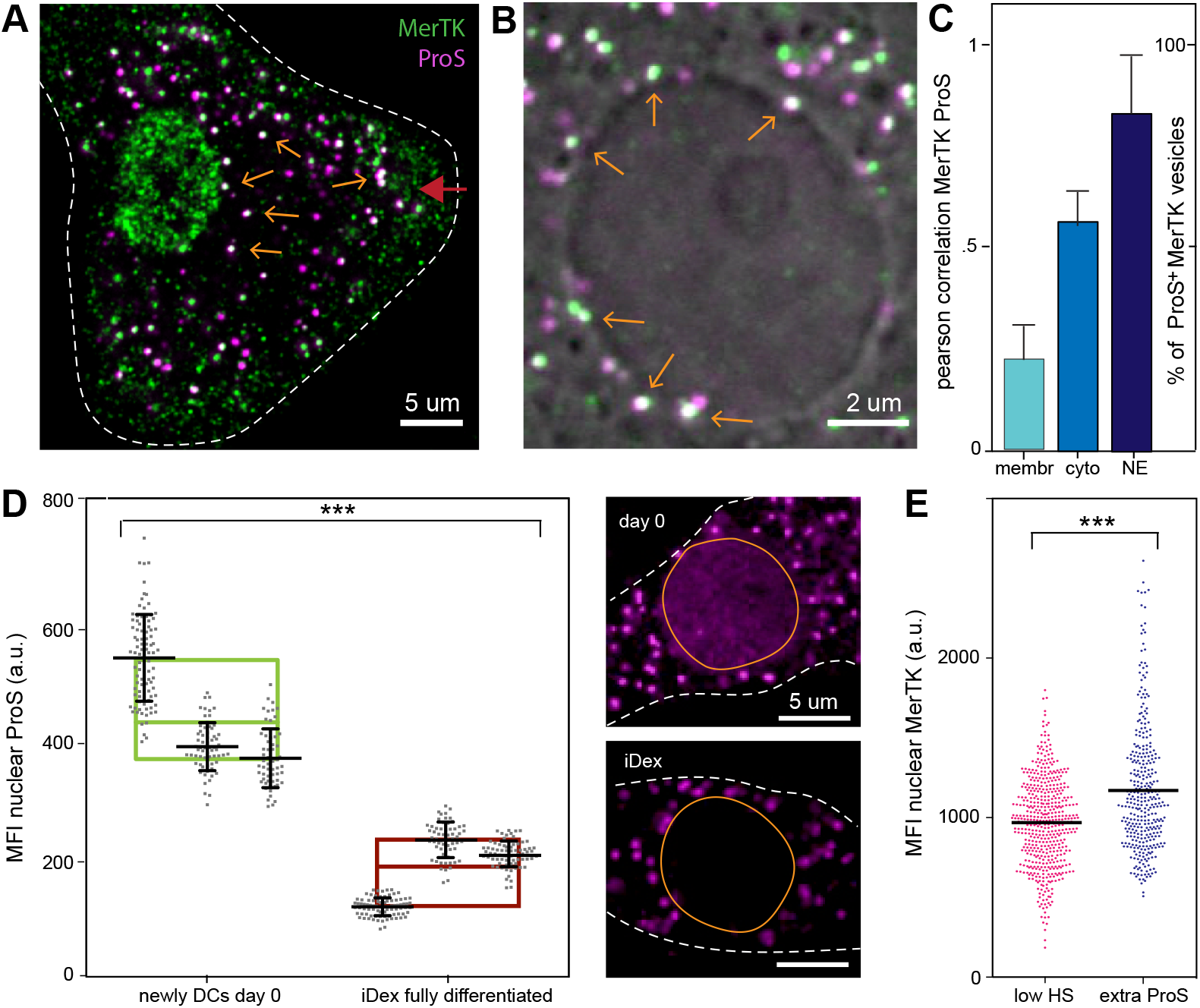
Effect of ligand ProS binding on translocation of *n*MerTK. (**A**) Representative dual colour confocal image of MerTK and its ligand ProS at day 0. MerTK is shown in green, ProS in magenta. Orange arrows indicate white spots in which both molecules clearly colocalize. The red arrow indicates the flattest part of the cell where the apical membrane is in focus. (**B**) Zoomed in on the nuclear area and overlaid with a transmission image to indicate the nucleus and its surrounding envelope. Orange arrows indicate spots of MerTK-ProS colocalization that are associated to the NE. Cells are minimally permeabilized in order to clearly observe the fluorescent spots at the NE. In these conditions there is minimal penetration of the antibodies into the nucleus. (**C**) Quantification of colocalization as a function of the cell region, i.e., membrane, cytoplasm and NE. Areas with the apical membrane in focus were chosen for the membrane portion of the analysis, the rest of the cell body excluding the nucleus was categorized as the cytoplasm. On zoom-in images like B, we manually counted the percentage of MerTK spots at the NE that colocalize with ProS. 10-20 cells from 2 different donors. (**D**) MFI of *n*ProS at day 0 and after full differentiation into iDexs. Each spot represents a single nucleus; the smaller plots correspond to 3 different donors measured. Side panels provide representative fluorescence images of ProS at these time points. (**E**) MFI of *n*MerTK with and without the addition of extra recombinant human ProS during culture. Data from 3 different donors.

To further demonstrate the involvement of ProS in *n*MerTK translocation we performed similar imaging experiments on newly differentiated DCs where *n*MerTK levels were found to be maximum (day 0, Fig. 2A). We hypothesized that more ProS would be found in the nucleus of the cells at day 0 as compared to day 4 (iDex cells, Fig. 3A-C). We found a 2.5-fold increase of nuclear ProS at day 0 compared to day 4 (Fig. 3D), supporting the notion that ProS indeed plays an important role in facilitating *n*MerTK trafficking. Since DCs require the presence of human serum (HS) in their growth medium and HS naturally contains high levels of ProS, it is not feasible to fully deprive the cells of ProS to further investigate its direct effect on *n*MerTK translocation. As an alternative, we cultured DCs in the presence of highly reduced HS levels, and compared *n*MerTK accumulation to cells cultured in the same reduced serum conditions but with the extra addition of soluble ProS. A significant increase in the amount of *n*MerTK was observed at these higher levels of ProS (Fig. 3E), further strengthening our findings that *n*MerTK translocation is ligand dependent.

### The endocytic receptor LRP-1 facilitates *n*MerTK translocation

We showed that MerTK expression is upregulated at two different stages during the differentiation of monocytes into iDex. First, at day 0, where upregulation is accompanied with a high localization of the receptor in the nucleus (Fig. 2A) and second, on fully differentiated iDex, where upregulated MerTK is mostly associated to the plasma membrane (Fig. 1A and Ref. 13). Although we showed that *n*MerTK translocation is facilitated by ProS, serum levels of ProS are constant through the entire DC differentiation process and as such, the ligand on its own is not likely to fully determine the spatial destination of the receptor. We thus hypothesized that MerTK requires an additional factor chaperoning its shuttling towards the nucleus at day 0 and that moreover, this factor must be lacking (or downregulated) in fully differentiated iDex, where MerTK remains largely membrane associated (Fig. 1A and Ref. 13). An interesting candidate for this differential spatial regulation is the endocytic receptor LRP-1, which is known to form a complex with Axl to facilitate internalization (Subramanian et al., 2014). Axl and MerTK are close relatives within the TAM family, making it conceivable that MerTK and LRP-1 can interact in a similar manner. Furthermore, LRP-1 has been reported to play a role in the shuttling of soluble environmental factors into the nucleus (Chaumet et al., 2015). To elucidate whether LRP-1 plays a shuttling role for MerTK, we performed intracellular dual colour confocal imaging of both MerTK and LRP-1. A remarkably strong colocalization between both receptors was observed (Fig. 4A) with multiple receptor complexes associated to the NE (orange arrows in Fig. 4B and quantification over multiple cells in Fig. 4C).

**Figure 4:**
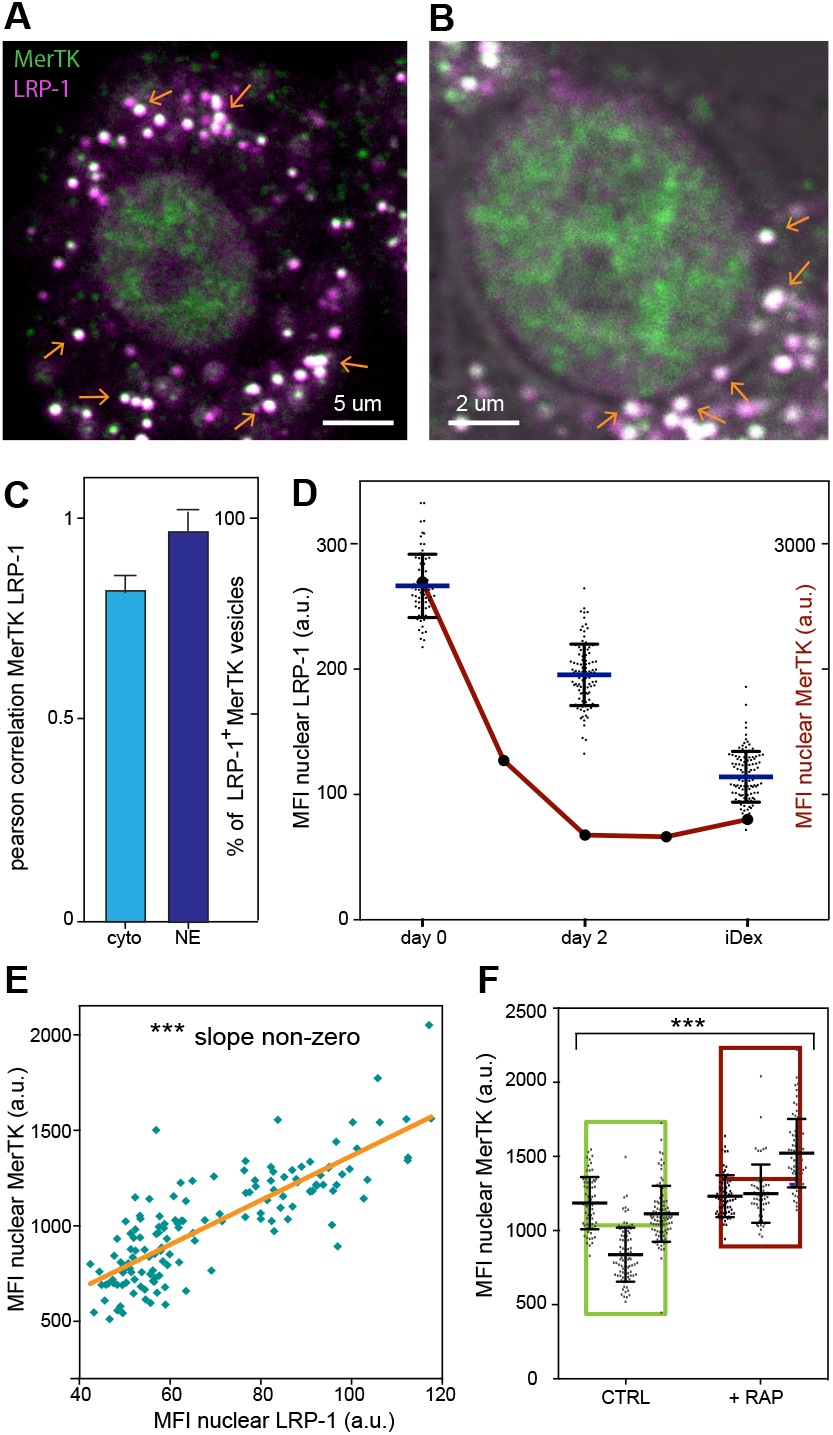
Role of LRP-1 in translocation of *n*MerTK. (**A**) Representative dual colour confocal image of MerTK (green) and LRP-1 (magenta) intracellularly. Orange arrows point to spots of clear colocalization between both receptors. (**B**) Representative zoomed in on the nuclear area, showing MerTK and LRP-1 signals overlaid with a transmission image (grey scale) showing the nucleus and NE. Orange arrows indicate spots in which MerTK and LRP-1 are colocalized and associated to the NE. (**C**) Quantification of the degree of colocalization between MerTK and LRP-1, both intracellularly and at the NE. Data from 3 different donors. (**D**) MFI of *n*LRP-1 over time in culture, for day 0, day 2 and iDexs (= day 4). The red curve together with the right y-axis corresponds to the same data of *n*MerTK shown in Fig. 2A, to facilitate comparison. Data from 3 donors. (**E**) Correlation between MFI of both *n*MerTK and *n*LRP-1 obtained from individual cells. Data from cells with and without the addition of RAP are included. (**F**) MFI of *n*MerTK with and without the addition of RAP to the culture (from day −2 to day 0). Each spot represents a single nucleus; small plots in larger bars represent data from 3 different donors.

We then quantified the amount of nuclear (*n*)LRP-1 as a function of DC differentiation. Interestingly, *n*LRP-1 expression follows a similar trend as to *n*MerTK, i.e., being highest at day 0, and decreasing steadily as a function of DC differentiation (Fig. 4D). However, unlike MerTK whose total expression increases again towards the final stage of differentiation (mostly located on the membrane), the total expression levels of LRP-1 remain low on fully differentiated iDex (Supplementary Fig. 4A). These results thus suggest that LRP-1 might play a role in the spatial partitioning of MerTK, determining whether nuclear translocation or membrane expression takes place.

In a model in which LRP-1 acts as a chaperone in bringing MerTK from the membrane to the nucleus, one would expect to find a positive correlation between the expression levels of both proteins in the nucleus. Taking advantage of naturally occurring cell to cell variability, we quantified the levels of *n*MerTK and *n*LRP-1 at the single cell level over multiple cells. Indeed, as suspected, a positive correlation between the translocation of both receptors was obtained (Fig. 4E). To inquire which receptor is leading and which receptor is following in this correlated translocation, we stimulated LRP-1 by adding one of its many ligands, RAP, in the medium. RAP-stimulation significantly increased nuclear LRP-1 translocation (Supplementary Fig. 4B) and most importantly, it also led to increased *n*MerTK translocation (Fig. 4F), again pointing towards a facilitating role of LRP-1 in the shuttling of MerTK into the nucleus. Overall, our results strongly suggest that LRP-1 plays a major role as chaperone molecule in regulating the sub-cellular spatial partitioning of MerTK, either to the nucleus or to the membrane.

### *n*MerTK is associated to chromatin, preferentially in an open conformation

The intriguing findings of the existence of a *n*MerTK population in immune cells prompted us to assess its specific nuclear location at the nanoscale, as well as its potential role. For this, we first separated day 0 DCs and DC-like THP-1 cells that both highly express *n*MerTK in different cellular fractions: the cytoplasm, the soluble part inside the nucleus, and the chromatin-bound fraction (Wang et al., 2015). These fractions were subsequently analysed by Western blotting with an antibody against MerTK. MerTK was found in all three fractions in both DCs and DC-like THP-1 cells, including the chromatin-bound fraction (Supplementary Fig 5A). This result suggests that *n*MerTK could play a role in gene expression regulation.

To investigate the spatial relationship between *n*MerTK and chromatin at the molecular level, we used dual colour Stochastic Optical Reconstruction Microscopy (STORM) following the approach of Ricci et al (Ricci et al., 2015) (Supplementary Fig 5B,C). This super-resolution technique allowed us to identify individual fluorescently-labelled *n*MerTK and histone molecules within the crowded environment of the nucleus with a localization precision of about 20 nm (Fig. 5A). In mammalian cells, the nuclear periphery is enriched in condensed heterochromatin, generally associated with transcriptional repression (Dekker and Misteli, 2015). Several studies have further demonstrated a direct link between the association of chromatin to the nuclear lamina and gene silencing (Finlan et al., 2008; Guelen et al., 2008; Kosak et al., 2002; Reddy et al., 2008). Consistent with these published results, our STORM images showed the condensed heterochromatin as a dense ring at the edge of the nucleus (Fig. 5A). Interestingly, this ring was mostly deprived of *n*MerTK (Fig. 5A, upper right panel). Remarkably, *n*MerTK was observed in the central nuclear region where the chromatin is much less dense (euchromatin) (Fig. 5A, lower right panel). In these regions, we also observed elongated structures composed of *n*MerTK and histones (Fig. 5A, pink dotted lines) that resemble a configuration where the nucleosomes are well-separated, DNA occupancy is low and chromatin is accessible (Ricci et al., 2015). The strong localization of *n*MerTK to nuclear regions where DNA is in an accessible configuration together with its clear exclusion from dense heterochromatin regions suggest that *n*MerTK might interact with active genomic regions.

**Figure 5:**
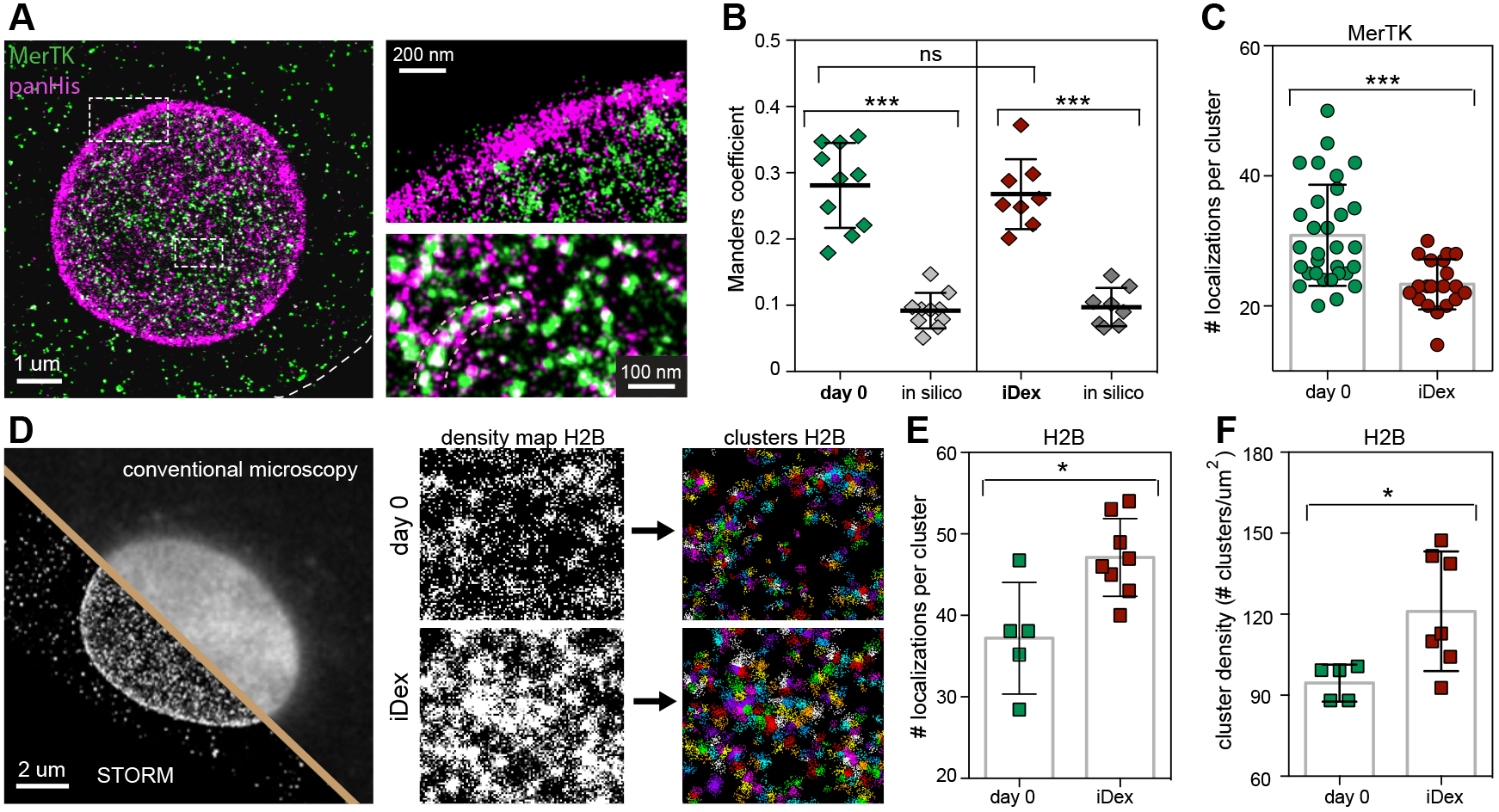
Dual colour super resolution STORM imaging of *n*MerTK and chromatin. (**A**) Representative reconstructed dual colour STORM image of *n*MerTK (green) and panHis (magenta) in the nucleus of a day 0 DC. The nucleus is delineated by a dense ring of histones (heterochomatin). The dotted white line indicates the cell boundary. The panels on the right correspond to zoom in images at two different nuclear areas. Elongated structures composed of *n*MerTK and histones are visible in the bottom zoom-in image (dotted pink lines). (**B**) Quantification of the colocalization between panHis and *n*MerTK (after image processing shown in Suppl. Fig. 5D) using the Manders overlap coefficient. The degree of colocalization was determined at day 0 and iDex, and was in both cases compared to the degree of colocalization between the experimental distribution of panHis and a random distribution of MerTK (*in silico*). Each symbol in the plot corresponds to an individual nucleus analysed. Data from 2 different donors. (**C**) Quantification of the number of localizations per *n*MerTK nanocluster in the nucleus of both day 0 and iDex DCs. Each dot corresponds an individual cell, with an averaged value from hundreds of nanoclusters per cell. Data from 3 different donors. (**D**) Representative single-color reconstructed STORM image of H2B in the nucleus of a day 0 DC. The image was partially overlaid with a conventional image of H2B in the same cell, to show the increase in resolution gained by using STORM. The panels on the right show the image processing strategy to generate density maps of the H2B signal (see methods for details). (**E**) Quantification of the number of localizations per H2B nanocluster in day 0 and iDex DCs. (**F**) Quantification of the H2B nanocluster density (the number of nanoclusters per μm^2^) in day 0 and iDex DCs. Each square in (E) and (F) corresponds to the average value over hundreds of H2B nanoclusters/cell of a single cell.

To further investigate this possibility, we quantified the degree of spatial association between *n*MerTK and euchromatin on manually selected central nuclear regions (excluding the nucleoli) in the STORM images (Supplementary Fig. 5D). In addition, we estimated the degree of random colocalization occurring as a result of the high density of histones and *n*MerTK by generating *in-silico* images of randomly distributed *n*MerTK molecules (using the experimentally obtained *n*MerTK density in that particular cell). We super-imposed the *in silico* data to histone STORM images and calculated the degree of random colocalization. A high degree of colocalization was found on the experimentally obtained STORM images, well-above random values and similar for day 0 DCs and for iDex cells (Fig. 5B). To further confirm that the observed colocalizations are real and not the result of cross-talk during imaging and/or cross-reactivity of the antibodies, we focused on areas where signal from only one of the two proteins is expected: cytosolic vesicles in the case of MerTK (Supplementary Fig. 5E) and the heterochromatin ring in the case of histones (Supplementary Fig. 5F). In both cases, the cross-talk was < 2%. These results thus confirm a high degree of spatial association between *n*MerTK and histones in central nuclear regions, strongly suggesting that *n*MerTK interacts with euchromatin in human DCs.

### Chromatin compaction increases upon DC differentiation and correlates with a reduction of *n*MerTK accumulation into nanoclusters

Surprisingly, the results in Fig. 5B showed a similar degree of association between *n*MerTK and euchromatin regardless of DC differentiation state, i.e., day 0 and iDex, whereas the confocal data revealed higher levels of *n*MerTK in day 0 than in fully differentiated iDex cells (Fig. 2A). To understand this apparent discrepancy, we quantified the nanoscale organization of *n*MerTK on both cell types from STORM images using a cluster analysis algorithm as described by Ricci et al (Ricci et al., 2015) (see materials and methods). A much larger number of localizations per nanocluster was observed at day 0 compared to iDex (Fig. 5C) whereas the total number of nanoclusters per area was similar on both cell types (Supplementary Fig. 5G). These results thus reveal a direct correlation between *n*MerTK levels and nanocluster size, rather than nanocluster density. Altogether, these data suggest that *n*MerTK nanoclusters might constitute functional units associated to chromatin that could more potently operate in newly differentiated DCs where *n*MerTK clusters are larger.

The results above naturally arise the question of how chromatin is organized on newly differentiated DCs (day 0) versus fully differentiated iDex (day 4). To address this question, we stained histone H2B in both cell types, performed STORM imaging over multiple cells (Fig. 5D) and compared their organization (see materials and methods). Interestingly, we found an increased number of H2B localizations per cluster (Fig. 5E) as well as increased cluster density (Fig. 5F) on fully differentiated iDex as compared to Day 0 DCs. The simultaneous increase in both parameters indicate that fully differentiated DCs have more histones covering their DNA and that chromatin is therefore more compact. We confirmed these results using conventional wide-field imaging, obtaining an increase of 30% in histone expression levels in the nucleus of iDex cells (Supplementary Fig. 6). These results are fully in line with a recent study showing a similar increase in the total histone content in pluripotent versus differentiated cells(Karnavas et al., 2014). An additional study also showed that histone content in monocytic-derived DCs can vary significantly upon treatment with different immunological stimuli (Parira et al., 2017). As H2B is directly involved in DNA compaction, our results reveal that chromatin is indeed in a more accessible conformation during early stages of DC differentiation, i.e., at day 0 DCs. Together, our observations that *n*MerTK shows increased clustering and preferentially interacts with chromatin exactly during this transcriptionally active stage where it is more accessible, together with its tendency to associate to euchromatin rather than to heterochromatin, strongly points towards a role for *n*MerTK as a genome regulator during DC differentiation.

### *n*MerTK has the potential to function as a transcription factor

A few recent reports have speculated on the possible function for RTKs in the nucleus, and proposed roles in DNA replication (Wang et al., 2006), repair (Liccardi et al., 2011) and/or transcription (Huo et al., 2010; Lin et al., 2001; Liu et al., 2010). Since DCs are non-proliferative cells, we conjectured that *n*MerTK could have a transcriptional function. Transcription factors are typically characterized by the presence of one or several transactivation domains that are involved in the recruitment of larger multiprotein complexes facilitating transcriptional activity (Raj and Attardi, 2017; Wärnmark et al., 2003). The sequence of these domains is highly conserved and can be predicted based on hydrophobicity and the presence of several key amino acids (Piskacek et al., 2016, 2007). We used an algorithm developed by Piskacek et al (Piskacek et al., 2007) to predict possible 9aaTAD (nine amino acid transactivation domain) regions in the MerTK sequence, and obtained two putative regions with a 100% match, within the cytoplasmic domain (Fig. 6A and Supplementary Fig. 7). This prompted us to test whether the cytoplasmic domain of MerTK indeed displays transactivational activity in a model cell system.

**Figure 6:**
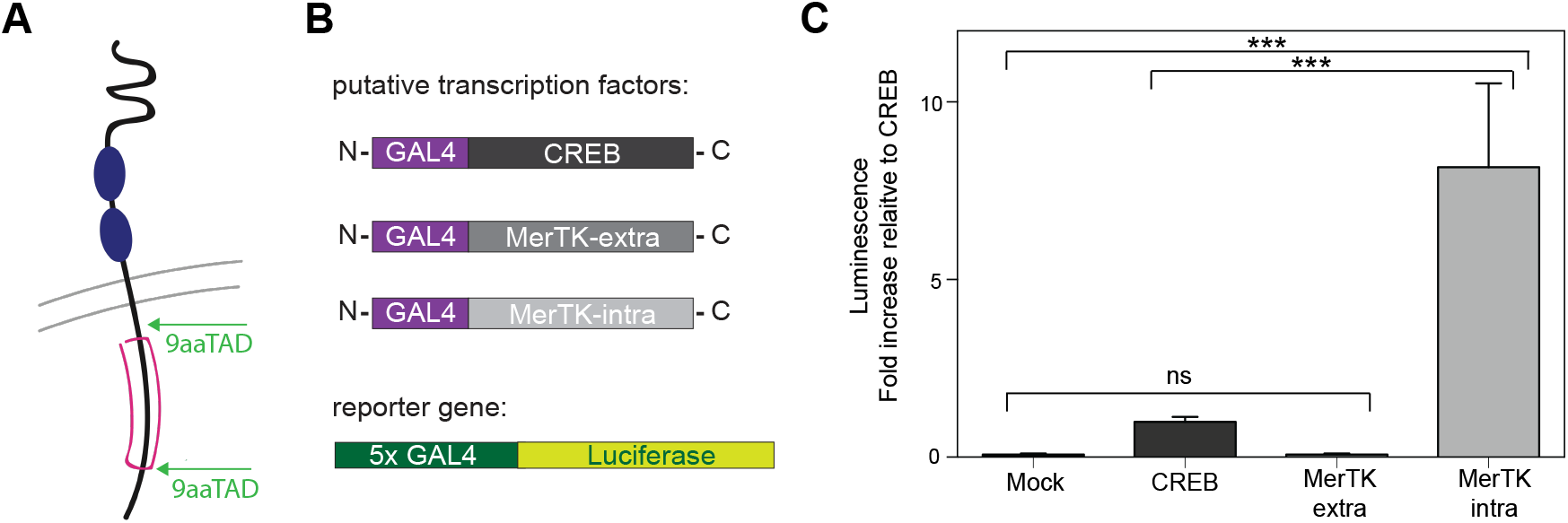
Transactivational activity of MerTK. (**A**) Schematic representation of MerTK topology (extracellular up) with predicted sites of 9aaTAD transactivation domains (both intracellular). (**B**) Schematic representation of the 3 fusion proteins that serve as putative transcription factors and the corresponding reporter gene. HeLa cells were co-transfected with one of the transcription factors and the reporter gene. (**C**) Quantification of the transcription induced by the different putative transcription factors, calculating the amount of luminescence generated after each co-transfection as described in (B), or Mock transfection as a negative control. Experiments were performed in triplicate and repeated 3 times during different cell line passages.

This intracellular domain, the extracellular domain and a positive control CREB (Sun et al., 1994), were fused to part of the DNA-binding protein GAL4, creating three different potential transcription factors (Fig. 6B). HeLa cells were co-transfected with a plasmid coding for one of these proteins, together with a reporter gene containing five GAL4 binding sites and a part coding for luciferase (Fig. 6B). The degree of luminescence found in the HeLa cells serves as a read-out for transcriptional activity induced by each of the possible transcription factors. As expected, the CREB fusion protein acted as a transcription factor and increased the transcription of the luciferase reporter gene compared to the control mock sample (Fig. 6C). The data was then normalized to the transcription induced by CREB, and displayed as a fold increase (Fig. 6C). Remarkably, the MerTK-intracellular fusion protein showed an enormous potential as transcription factor, with an eight-fold increase in transcription as compared to CREB (Fig. 6C). The MerTK extracellular domain did not induce any transcription beyond that of the Mock control, fully in line with the prediction that only the intracellular domain contains 9aaTAD sites and therefore transactivational capacity. Our data thus shows for the first time that MerTK has the potential to act as a potent transcription factor, strongly suggesting that *n*MerTK found in human DCs and associated to chromatin has the function to regulate gene transcription. Considering the time-sensitive dependence of nuclear accumulation of MerTK on DC differentiation, this transcription factor is likely to boost the upregulation of key genes during this process.

## Discussion

In this study we have identified for the first time, to our knowledge, the presence of the transmembrane receptor MerTK in the nucleus of human DCs and show that the degree of nuclear localization strictly depends on DC maturation. Our super-resolution STORM studies on intact nuclei further revealed that *n*MerTK preferentially associates to open and active chromatin. We found that chromatin compaction increases upon DC differentiation and correlates with a reduction of *n*MerTK accumulation into nanoclusters. We also showed that MerTK has the potential to act as a powerful transcription factor, suggesting that this transmembrane receptor regulates the expression of key genes during DC differentiation.

Although MerTK has previously been observed in the nucleus of a leukaemia cell line (Jurkat T cells) (Migdall-Wilson et al., 2012), our study is the first to give functional importance to this intracellular localization in the context of immunity in primary human cells. Based on our spatial, temporal and functional data, we suggest that *n*MerTK acts as a transcription factor involved in regulating the differentiation of human DCs in a time-sensitive manner. Interestingly, MerTK has been proposed to regulate the differentiation of natural killer (NK) cells by upregulating the membrane expression of certain key NK cell immunoreceptors during a well-defined period of cell maturation (Sun et al., 1994). The authors of that study envisioned that such upregulation happens through classical downstream signalling and activation of traditional transcription factors. However, they found that the upregulation was not caused by any of the known transcription factors involved in NK cell development. In the light of our work, it is highly conceivable that just like in DCs, MerTK translocates to the nucleus of NK cells to induce the upregulation of several important immunoreceptors, thereby regulating differentiation in a time-sensitive manner. A previous study by Schmahl et al. indeed suggested a similar regulating role for the FGF2a-RTK in the early stages of Sertoli cell differentiation (Schmahl, 2004). Further studies focusing on the genes that MerTK regulates as a transcription factor are necessary to fully understand how *n*MerTK directs DC differentiation. We persistently attempted to perform ChIP-Seq profiling experiments to identify genomic regions influenced by MerTK, as well as nuclear IP of MerTK followed by quantitative mass spectroscopy to detect the nuclear factors that MerTK forms a complex with. Unfortunately, these experiments turned out unfeasible due to the lack of validated MerTK antibodies for these techniques together with the enormous demand of cellular material that is incompatible with the isolation and culturing of monocyte derived DCs from blood.

Our super-resolution studies on tolerogenic DCs showed the presence of small MerTK nanoclusters on the cell membrane. Although receptor dimerization is expected to occur as a result of ligand activation by ProS in the medium, our observation of more extensive nanoclustering is an important finding in the field of immunoreceptor organization at the cell surface. Membrane-bound MerTK expressed by tolerogenic DCs is thought to suppress the T cell response by depriving the local environment of ProS, a T cell activating factor (Cabezón et al., 2015; Carrera Silva et al., 2013). Efficient ProS scavenging from the local environment by MerTK requires rapid internalization of MerTK-ProS complexes in order to interfere with T cell binding and activation. We thus speculate that MerTK nanoclustering might provide an advantage for this rapid internalization by lowering the amount of energy and resources needed for ProS clearance. In addition, it is also conceivable that MerTK nanoclustering increases the binding affinity to ProS, favouring internalization. Moreover, the small number of MerTK molecules involved in each nanocluster (on average 3, ranging between 1 and 11) would provide an excellent strategy to optimize MerTK resources for efficient ligand scavenging throughout the local cell environment.

LRP-1 has been described as a receptor that regulates the protein composition of the plasma membrane (Gonias et al., 2004). In the context of immunity, LRP-1 regulates the membrane levels of β1 integrins (Theret et al., 2017; Wujak et al., 2018), CD44 (Perrot et al., 2012) and the phagocytic receptor AXL (Subramanian et al., 2014) by facilitating their endocytosis. Knock-down or blocking of LRP-1 leads in all cases to an accumulation of the receptors at the membrane level. Our results on the role of LRP-1 in the partitioning and regulation of the spatial location of MerTK can be fully rationalized under the paradigm that LRP-1 controls the composition of the cell membrane. Like the previously mentioned receptors, we found that MerTK accumulates at the cell membrane when LRP-1 expression levels are low, which physiologically occurs during the final stages of DC differentiation (day 4). However, while LRP-1 targets many other receptors towards lysosomal degradation or recycling, we show here that this receptor is also involved in nuclear translocation. It was shown previously that LRP-1 can target soluble toxins from the cell environment into the nucleus in a receptor-ligand fashion (Chaumet et al., 2015), but to our knowledge LRP-1 has never been implicated in chaperoning other transmembrane proteins towards the nucleus. We thus propose that the subcellular spatial destination of MerTK is tuned by LRP-1. When DCs simultaneously express both MerTK and LRP-1, LRP-1 will bind MerTK and will direct it into the nucleus (day 0) via endocytosis. By contrast, in the absence of, or at reduced LRP-1 levels, assistance for nuclear translocation is compromised, and MerTK remains at the surface (day 4). Pinpointing, for the first time, the role of LRP-1 in this process is a great step forward in the molecular understanding of nuclear trafficking of transmembrane receptors. This knowledge can be used to further our understanding of nuclear translocation of other RTKs. Since many other RTKs play an oncogenic role in the nucleus, identifying triggers in this process is of paramount importance for future clinical interference.

The presence of full-length RTKs in the nucleus has been reported for a significant number of receptors over the last decade (reviewed in (Brand et al., 2012; Wang and Hung, 2012; Wells and Marti, 2002a)). However, the community has been rather reluctant to accept these observations, partially because it is counter-intuitive to envision how full-length membrane-bound receptors containing a hydrophobic transmembrane domain could be soluble inside the nucleoplasm. The existence of RTKs with a deleted transmembrane domain has been proposed in a model in which those mutated proteins dimerize with their wild-type counterparts (Wells and Marti, 2002a). The dimerization would provide ligand sensitivity and explain its localization at the cell membrane as well as the membrane of intracellular compartments. Such a soluble, almost full-length isoform has been detected for the FGFR2 receptor (Katoh et al., 1992), but for many other RTKs in the nucleus, including MerTK, the existence of such an isoform has never been demonstrated. Moreover, the specific presence of the transmembrane domain in the nucleus has been shown in some cases (Wells and Marti, 2002b), contradicting this model. Alternatively, we hypothesize that although not incorporated in the lipid bilayer, the transmembrane domain could still be covered by a small amount of lipids in the nucleus. The concept of nuclear lipids has been widely described over the past two decades, and the existence of proteolipid complexes has been observed (Albi and Magni, 2004; Cascianelli et al., 2008; Irvine, 2000). *In vitro* experiments along these lines will be important to further understand the intriguing phenomenon of soluble transmembrane receptors as nuclear regulators.

A second aspect discouraging the investigation of nuclear localization of membrane proteins is the general consensus that aberrant or overexpression of the protein causes this atypical nuclear translocation, and that the presence of RTKs in the nucleus is mostly related to malignancies. Our results on MerTK are however markedly distinct in several ways: *First*, *n*MerTK is found in a very high concentration in healthy non-proliferating DCs. *Second*, *n*MerTK localization seems to be exclusively reserved for immune cells (directly isolated DCs, monocyte-derived DCs and THP-1 cells in our experiments, and Jurkat T cells (Migdall-Wilson et al., 2012)). Indeed, we showed that overexpression of MerTK in other cell types, both with (HEK293) and without (HeLa) endogenous MerTK, does not lead to nuclear accumulation of the receptor. *Third*, the degree of nuclear translocation strictly relates to DC differentiation, a physiological process that is crucial to immunity. The sharp peak of *n*MerTK accumulation we observed in newly differentiated DCs suggests a critical time-sensitive and well-regulated need for the presence of the receptor in the nucleus during differentiation. Our results thus interestingly suggest a physiological, non-malignant role for a RTK in the nucleus, validating the importance of further studies on this puzzling and unconventional way of cell signalling.

## Materials and methods

### Cell culture

Dendritic cells were derived, as reported previously(de Vries et al., 2005), from peripheral blood samples. Buffy coats from healthy donors were obtained from *Banc de Sang i Teixits* upon written informed consent. In brief, peripheral blood mononuclear cells (PBMCs) were allowed to adhere to a plastic surface for 2 h at 37°C. Unbound PBMCs were washed away, and the remaining adherent monocytes were cultured for 48h in the presence of IL-4 (300 U/ml) and GM-CSF (450 U/ml) (both from Miltenyi Biotec, Madrid, Spain) in X-VIVO 15 (BioWhittaker, Lonza Belgium) medium supplemented with 2% AB human serum (Sigma-Aldrich, Spain). At that moment, they are day 0 DCs, and were used for several experiments. To generate iDexs, the cells were further cultured for 4 days in the same conditions plus Dexamethasone (1 μM; MERCK). IDCs were equally generated in 4 days, but without the extra addition of Dex. For serum starvation experiments, monocytes were cultured normally up to day 0. Then, they were differentiated into iDex DCs using cytokines and Dex, but using a lower concentration of HS (1% instead of 10%). This percentage was experimentally determined as the lowest concentration at which the DCs still developed normally (assessed visually). During the last 48h of differentiation, recombinant human ProS was added to one of the conditions (concentration according to Cabezón et al. (Cabezón et al., 2015)).

THP-1 cells were cultured in RPMI 1640 medium supplemented with antibiotic-antimycotic (both Gibco) and 10% FBS (ThermoFisher). To induce a DC-like phenotype, they were cultured for 6 days in the presence of IL-4 and GM-CSF, with a medium exchange after 3 days.

HeLa cells and Hek293 cells were both cultured in complete medium (Dulbecco’s modified Eagle’s medium containing 10% fetal bovine serum (both Gibco)).

### Antibodies and reagents

The following primary antibodies were used throughout this study at a concentration of 5 μg/ml, except for the STORM experiments where they were used at a concentration of 20 μg/ml: α-MerTK (mouse extracellular monoclonal, 125618, R&D Systems), α-MerTK (goat extracellular polyclonal, AF891, R&D Systems), α-MerTK (rabbit extracellular monoclonal, Y323, Abcam), α-MerTK (rabbit intracellular polyclonal phosphospecific, PMKT-14GAP, FabGennix), α-ProS-AF647 (bs-9512R-A647, Bioss), α-LAMP1 (H5G11, Santa Cruz Biotechnology), α-calreticulin (ADI-SPA-601, Enzo), α-EEA1 (14/EEA1, BD Biosciences), α-LRP-1 (LRP1-11, Sigma-Aldrich), α-PanHis (H11-4, Merck Millipore), α-H2B (5HH2-2A8, Merck Millipore), α-HDAC1 (10E2, Cell Signalling), α-tubulin (YL1/2, Abcam).

For confocal and STED imaging, the following secondary antibodies were used, all at a concentration of 10ug/ml: Goat-α-mouse-AF488 (A11001, ThermofFisher), Goat-α-mouse-Atto647N (50185, Sigma-Aldrich), Goat-α-rabbit-AF488 (A11008, ThermoFisher), Goat-α-rabbit-AF647 (A21244, ThermoFisher).

For Western blot imaging, the following secondary antibodies were used (all from ThermoFisher): Donkey-α-rabbit-AF680 for MerTK, Donkey-α-mouse-AF680 for HDAC1 Donkey-α-rat-DyLight800 for tubulin.

For STORM imaging, the secondary antibodies (donkey-α-mouse and donkey-α-rabbit from ImmunoResearch, used at a concentration of 20 μg/ml) were labelled in-house with different combinations of pairs of activator/ reporter dyes. The dyes were purchased as NHS ester derivatives: Alexa Fluor 405 Carboxylic Acid Succinimidyl Ester (Invitrogen), Cy3 mono-Reactive Dye Pack (GE HealthCare), and Alexa Fluor 647 Carboxylic Acid succinimidyl Ester (Invitrogen). Antibody labelling reactions were performed by incubating a mixture of secondary antibody, NaHCO3, and the appropriate pair of activator/reporter dyes diluted in DMSO for 40 min at RT. Purification of labelled antibodies was performed using NAP5 Columns (GE HealthCare). The dye to antibody ratio was quantified using Nanodrop and only antibodies with a composition of 3-4 Alexa Fluor 405 and 0.9-1.2 Alexa Fluor 647 per antibody were used for imaging.

Recombinant human PROS1 (R&D systems, 50 nM final concentration) and recombinant human RAP (Merck Millipore, 200 nM final concentration) were used to stimulate nuclear translocation of MerTK.

### Flow cytometry

For flow cytometry analysis, DCs were labelled with primary antibody α-MerTK (R&D systems), followed by secondary staining with PE-labelled goat-anti-mouse (from BD Biosciences), both for 30 min at 4°C and a concentration of 5 μg/ml. Appropriate isotype control IgG1 (from BD Biosciences), was included. Flow cytometry was performed using FACSCanto II.

### Plasmids

The MerTK (Mer cDNA ORF Clone, Human, untagged, pCMV3) was obtained from Sino Biological. Both pcDNAI-GAL4-CREB and 5xGAL4-TATA-luciferase were a gift from Richard Maurer(Sun et al., 1994) (Addgene plasmid # 46769 and Addgene plasmid # 46756, respectively). For transfection experiments of MerTK in different cancer cell lines, a GFP-tag was added to the plasmid. For the luciferase assay, we cloned the pcDNAI-GAL4-MerTK-extracellular and the pcDNAI-GAL4-MerTK-intracellular constructs by Gibson assembly of two fragments, the first one obtained by digesting the pcDNAI-GAL4-CREB vector with EcoRI and XbaI restriction enzymes (New England Biolabs), and the second part obtained by amplifying either the extracellular or the intracellular coding regions of the MerTK vector by PCR. The primers used for these amplifications (obtained from Integrated DNA technologies) were 5’-AGTAGTAACAAAGGTCAAAGACAGTTGACTGTATCGCCGGAATTCGCTATCACTGAGGCAAGGGAAGAAG-3’ and 5’GATCCTCTAGCATTTAGGTGACACTATAGAATAGGGCCCTCTAGAGATGATGAGCACAGGATCTTAGTT-3’ for the extracellular domain of MerTK (residues 21-505) and 5’-AGTAGTAA CAAAGGTCAAAGACAGTTGACTGTATCGCCGGAATTCAAAAGAGTCCAGGAGACAAAGTTTGG-3’ and 5’-GATCCTCTAGCATTTAGGTGACACTATAGAATAGGGCCCTCTAGATTACATCAGGACTTCTGAGCCTTCTGAGGAGT-3’ for the cytosolic domain of MerTK (residues 527-999). SnapGene software (obtained from GSL Biotech) was used for molecular cloning procedures.

### MerTK transfection

HeLa cells were transfected using TransIT-HeLaMONSTER and HEK293 cells using TransIT-293 (both from Mirus). Cells after transfection were cultured both with and without ProS or HS in the medium, to potentiate nuclear translocation of MerTK. Cells were typically imaged 24h after transfection, although both earlier and later time points were also explored.

### Transcriptional activation luciferase assay

HeLa cells were seeded on a 24-well plate, 2.5×10^4^ cells per well. After 24 hours, cells were cotransfected with both the reporter gene and one of the different putative transcription factors using X-tremeGENE 9 (Roche) following the manufacturer’s recommendations. The cells received 1 unit of 5xGAL4-TATA-luciferase reporter DNA and 0.4 units of the putative transcription factor DNA. 48 hours after transfection, cells were lysed with 100 μl of cell culture lysis reagent (Promega, Luciferase Assay System Kit #E1500) for 10 min on ice, and then spun down at 12000 g for 2 min at 4°C. For the luciferase assay we mixed 20 μl of those supernatants with 100 μl of the luciferase assay reagent (Promega, Luciferase Assay System Kit #E1500) in a well of a white, opaque 96-well plate, and the luminescence was measured after 30 sec using a manual luminometer (Gen5 microplate reader, BioTek), programmed to perform a 10 sec measurement read for luciferase activity. Luminescence was normalized and represented as the fold increase relative to the luminescence induced by positive control fusion protein GAL4-CREB. Each transfection was performed in triplicates, and the experiments were repeated 3 times in different days.

### Cell fractionation and Western blot detection

Dendritic cells at day 0 and DC-like THP-1 cells were collected and fractionated into the cytoplasmic fraction, the soluble nuclear fraction and the chromatin bound fraction following Wang et al.(Wang et al., 2015). In brief, cells were crushed using a Dounce tissue grinder set (Sigma-Aldrich) of 2ml, which homogenizes the cells without rupturing the nuclear membrane. Effectivity of this step was checked under the microscope with a Trypan Blue staining. Cytoplasmic material was then separated from the intact nuclei by centrifugation. The nuclei were subsequently lysed, and the chromatin was separated from the soluble fraction by centrifugation. The chromatin pellet was then sonicated in order to release associated proteins and allow their detection. All 3 fractions were then loaded and ran, transferred, and stained following standard Western blotting procedure. Besides MerTK, we also stained for tubulin and HDAC1 to verify the effectiveness of the cell fractionation.

### Sample preparation for fluorescence imaging

Fresh cells were diluted up to a concentration of 1×10^6^ per ml in plain medium, and attached to the bottom of the cover glasses (Lab-Tek) by incubation for 30 min. Samples were then fixed using 4% paraformaldehyde (PFA) for 15 min at RT. Then, cells were blocked and permeabilized for 1h at RT with 3% BSA and 0,5% TritonX-100 in PBS, followed by primary and secondary labelling both for 30 min at RT. Finally, all samples were fixed again with 2% PFA and stored at 4 °C. For membrane staining, TritonX-100 was left out from the blocking mixture.

### Confocal imaging

Imaging was performed using a confocal microscope (TCS SP5, Leica Microsystems). Images were taken with a 1.4 NA oil immersion objective (HCX PL APO CS 63.0x, Leica), a 512×512 pixels format and a scanning speed of 400 Hz. AF488 was excited with the 488 nm line, at 25% of the argon laser power and detected between 500 nm and 570 nm. Atto647N or AF647 was excited with the 633 nm line at 30% of the HeNe laser power and detected between 645 nm and 715 nm. To be able to use the fluorescence intensity measurements in a quantitative way, imaging conditions were always kept constant across measurements, and a calibration sample was used to account for day to day fluctuations in the system.

### STED imaging

Imaging was performed using a commercial STED microscope (TCS SP5, Leica Microsystems). Images were taken with a 1.4 NA oil immersion objective (HCX PL APO CS 63.0x, Leica), a 1024×1024 pixels format and a scanning speed of 1400 Hz. The effective imaging beam consisted of the 488 nm line, at 25% of the argon laser power, and 100% of the depletion donut-shaped laser at 592 nm. Fluorescence was collected between 500 nm and 570 nm.

### STORM imaging

Imaging was performed using a commercial microscope system from Nikon Instruments (NSTORM). Samples were prepared as described above, and imaged in the following buffer to facilitate blinking: Cysteamine MEA (Sigma-Aldrich), Glox Solution (0.5 mg/ml glucose oxidase, 40 μg/ml catalase; both Sigma Aldrich) and 10% Glucose in PBS(Bates et al., 2007). Images were acquired with a frame rate of 83 frames per second. In single color experiments, H2B was stained with the AF405-AF647 activator/reporter dye pair. By exciting AF405 with the corresponding laser line at 405 nm, this dye becomes activated and transfers its photons to the reporter dye. The reporter dye in turn will emit these photons only upon excitation with a 647 nm laser, and subsequently goes back into the dark state. We therefore used an imaging cycle in which one frame belonging to the activating light pulse (405 nm) was alternated with 3 frames belonging to the imaging light pulse (647 nm). Dual colour imaging was performed with two sets of secondary antibodies labelled with the same reporter dye (Alexa Fluor 647) but two different activator dyes (Alexa Fluor 405 for MerTK and Cy3 for panHis)(Bates et al., 2007). In addition to the first imaging cycle of 4 frames, a second cycle of 4 frames with an activation laser pulse at 561 nm was used to image Alexa Fluor 647 linked to activator Cy3.

In order to exhaustively image all fluorophores in a reproducible manner allowing for quantitative comparison across cells and conditions, we used the following scheme to increase the activator laser power, according to Ricci et al. (Ricci et al., 2015).

| Activating laser power | frames |
| --- | --- |
| 8 mW | Until 20.000 |
| 9 mW | 20.000 - 37.000 |
| 10 mW | 37.000 - 50.000 |
| 11 mW | 50.000 - 57.000 |
| 13 mW | 57.000 - 63.000 |
| 15 mW | 63.000 - 70.000 |
| 18 mW | 70.000 - 75.000 |
| 22 mW | 75.000 - 80.000 |
| 29 mW | 80.000 - 85.000 |
| 38 mW | 85.000 - 90.000 |
| 48 mW | 90.000 - 95.000 |
| 60 mW | 95.000 - 100.000 |

### STORM image reconstruction

STORM images were processed and rendered as previously described (Bates et al., 2007). Briefly, spots in single-molecule images were identified based on a threshold and fit to a Gaussian to identify their position in x and y. Applying this approach over all 100.000 frames gives the raw STORM data, consisting of a list of x-y coordinates, corresponding to the localized positions of all the fluorophores. Reconstructed images from the x-y coordinates were displayed using Insight3, after both drift and crosstalk correction following Refs 30 and 31.

Grouping of the x-y localization into clusters was done according to Ricci et al. (Ricci et al., 2015) using a custom-made cluster analysis algorithm written in MatLab. First, a density map was generated, in which each pixel has a value equal to the number of localizations falling within the pixel area (pixel size = 10 nm). A constant threshold was then used to convert the density maps into binary images, such that pixels have a value of 1 where the density is larger than the threshold and a value of 0 elsewhere. Localizations falling on zero-valued pixels were discarded from further analysis. For our threshold setting, the number of discarded localizations typically corresponded to < 5% of the total number of localization within a nuclear region. Connected components of the binary image, composed by adjacent non-zero pixels (4-connected neighbours), were sequentially singled out and analysed. Localization coordinates within each connected component were grouped by means of a distance-based clustering algorithm. Initialization values for the number of clusters and the relative centroid coordinates were obtained from local maxima of the density map within the connected region, calculated by means of a peak finding routine. Localizations were associated to clusters based on their proximity to cluster centroids. New cluster centroid coordinates were iteratively calculated as the average of localization coordinates belonging to the same cluster. The procedure was iterated until convergence of the sum of the squared distances between localizations and the associated cluster and provided cluster centroid positions and number of localizations per cluster.

### Image analysis

All image analysis was performed using ImageJ unless otherwise stated. Nuclear MFI (mean fluorescence intensity) from confocal images was quantified by manually selecting the nuclear area based on the transmission images of the cells. Colocalization was determined using the plugin Coloc2, and quantified by using the Pearson correlation coefficient for raw images, or the Mander’s overlap coefficient for binary images (in the case of the dual colour STORM images). Image segmentation was performed according to Rizk et al (Rizk et al., 2014), using their plugin. STED images were analysed using a custom-written routine in MatLab. From these images, the number of MerTK molecules per nanocluster on the membrane was estimated by dividing the background-corrected fluorescence intensity of each MerTK spot by the average intensity of the spots on the glass (single labelling units of 1 primary and several secondary antibodies). The physical size of the clusters was calculated by taking the FWHM (full width half max) of the fitting of the fluorescence intensity profile of each spot. The size of the spots on the glass provide the effective resolution of our STED system, around 100 nm.

### Statistical Analysis

All analyses were performed using GraphPad Prism 6. Results are shown as the mean ± SD. To determine statistical differences between the mean of two data sets, the (un)paired two-tailed Student T-test was used. To determine statistical differences between the mean of 3 or more data sets, the One-way ANOVA was used, followed by the Tukey’s multiple comparison test. On single-cell data coming from different donors, an average value per donor was used to calculate statistical differences. Significance is represented using: *ns (*P>0.05); * (P<0.05); ** (P<0.001) and *** (P<0.0001).

## Acknowledgements

The research leading to these results has received funding from the European Commission H2020 Program under grant agreement ERC Adv788546 (NANO-MEMEC), Spanish Ministry of Economy and Competitiveness (“Severo Ochoa” program for Centres of Excellence in R&D [SEV-2015-0522], FIS2015-63550-R (to M.K), FIS2017-89560-R (to M.F.G.-P.), BES-2013-064971 (to K.J.E.B), BFU2015-73288-JIN (to F.C), RYC-2017-22227-AEI/FEDER/UE (to F.C.), RYC-2015-17896 (to C.M.), BFU2017-85693-R (to C.M.), Fundació CELLEX (Barcelona), Fundació Mir-Puig and the Generalitat de Catalunya through the CERCA program and AGAUR (Grants No. 2017SGR940 to C.M. and No.2017SGR1000 to M.F.G.-P.).

## Author contributions

K.J.E.B., F.C., M.F.G.-P. designed the research. K.J.E.B., G.F.-G., and F.C. performed the experiments. K.J.E.B. and M.A.R performed STORM imaging. C.M. developed STORM analysis algorithms and provided advice on the analysis. M.L., A.C., D.B.-R., discussed data and hypotheses. K.J.E.B. and M.F.G.-P. wrote the manuscript. All the authors reviewed the manuscript and provided feedback.

## Supplementary Figures

**Figure S1:**
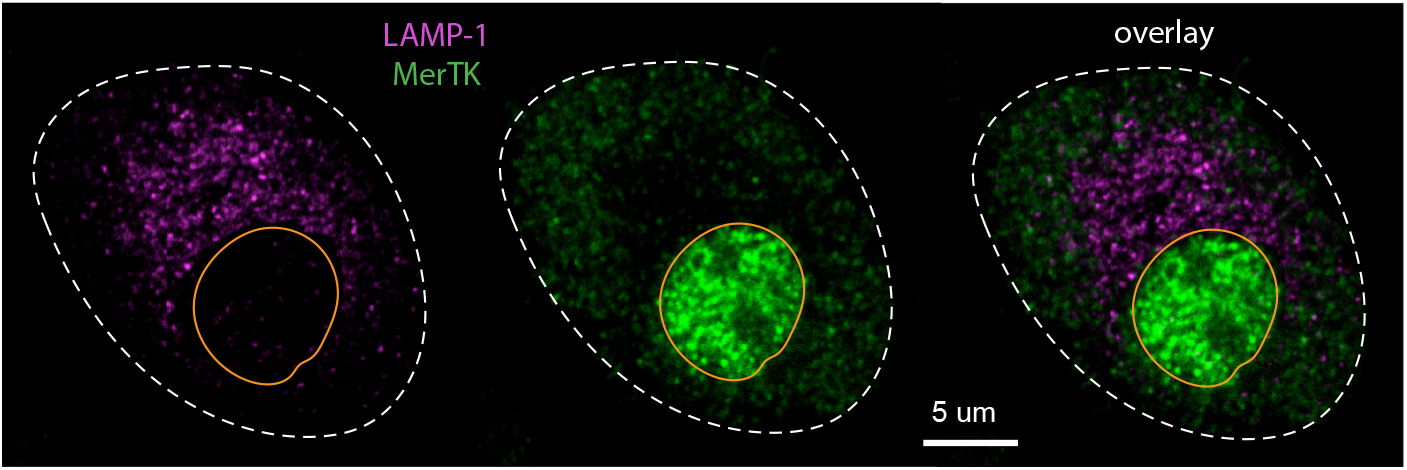
Intracellular MerTK does not reside in lysosomes. Representative dual colour confocal image of LAMP-1 staining the lysosomes in magenta, and MerTK in green. There is clear antilocalization visible between both components. The dotted line indicates the cell boundary, and the orange line the nuclear envelope.

**Figure S2:**
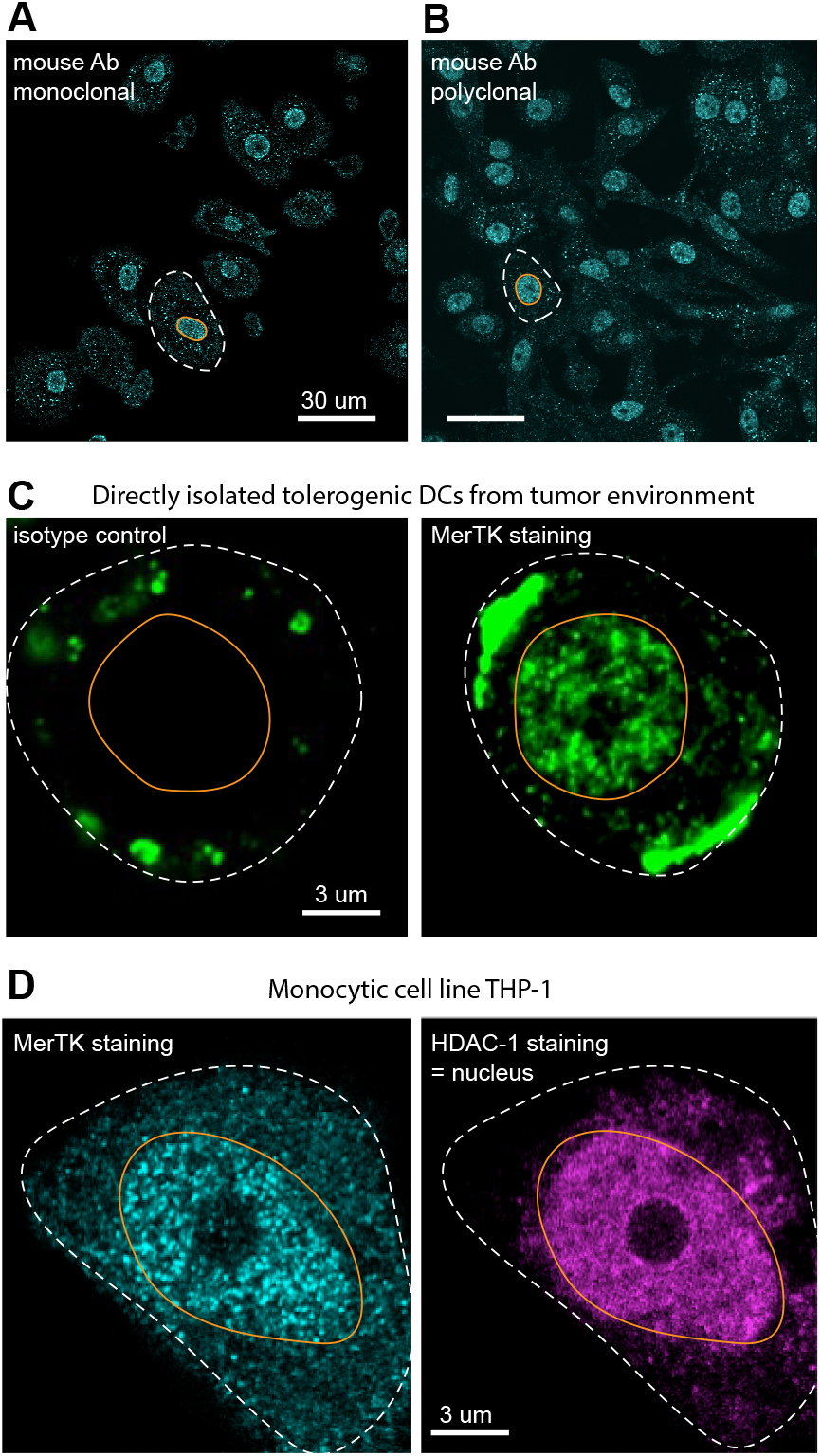
Nuclear staining of MerTK in different monocytic immune cells. (**A**) Representative confocal image of permeabilized iDex DCs stained with a mouse monoclonal antibody against an extracellular epitope of MerTK. The dotted line represents the cell boundary, while the orange line represents the nuclear envelope. (**B**) Representative image of permeabilized iDex DCs stained with a mouse polyclonal antibody against the extracellular domain of MerTK. (**C**) Representative confocal image of a permeabilized tolerogenic DC directly isolated from the tumor environment of a cancer patient. The cell is stained for MerTK, and nuclear localization becomes apparent, while this is not the case for the isotype control. (**D**) Representative confocal image of a permeabilized THP-1 cell (monocytic cell line) stained for MerTK. HDAC-1 staining is used to identify the nuclear area, as it less evident from the transmission images in the case of rounded monocytes.

**Figure S3:**
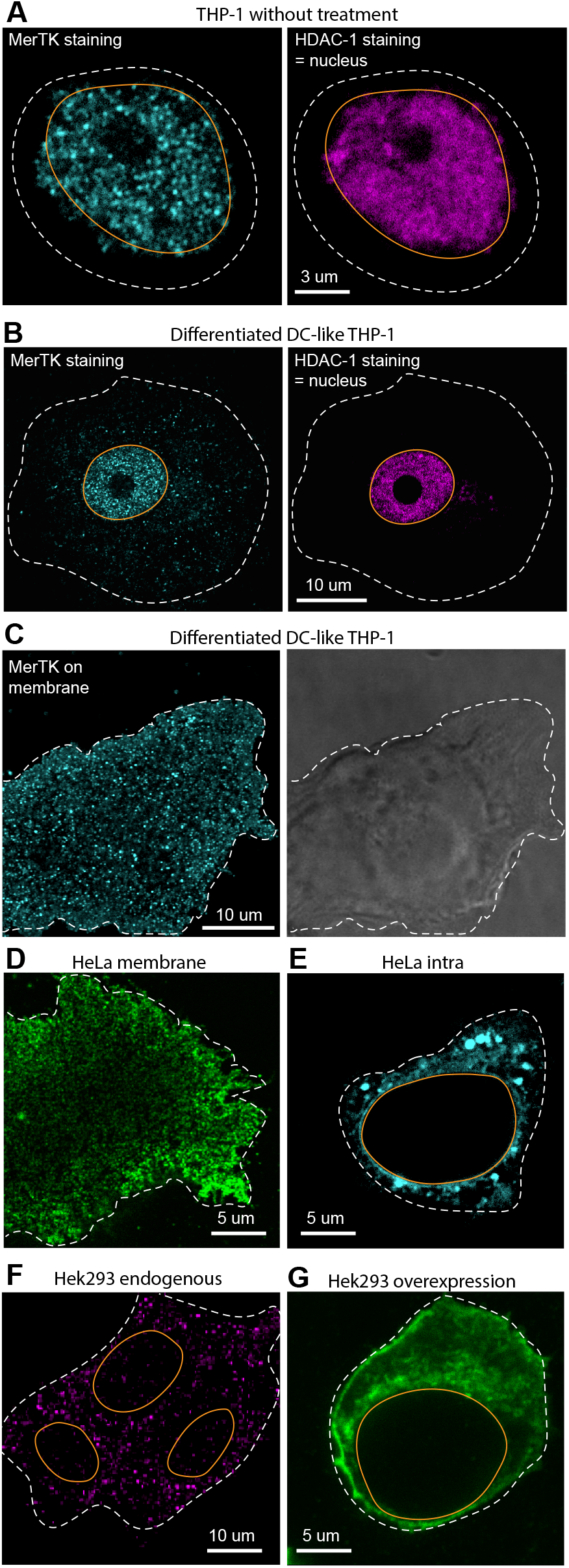
Nuclear localization of MerTK in immune cell lines vs other tissue cell lines. **(A)** Representative confocal image of a perm-eabilized THP-1 cell (monocytic cell line) stained for MerTK. The nuclear area is identified using a HDAC-1 staining, since it is less clear from transmission images for the rounded monocytes. The dotted line represents the cell boundary; the orange line represents the nuclear envelope. (**B**) Representative image of a permeabilized THP-1 cell that has been differentiated into a DC-like phenotype, stained for MerTK. The nuclear area is identified using HDAC-1 staining. (**C**) Representative image of a DC-like THP-1 stained for MerTK on the membrane. Nanoclustering of the receptor similar to that observed on the membrane of iDexs becomes apparent, as well as flattening and spreading of the cell. (**D**) Representative image of a MerTK transfected HeLa cell stained for MerTK on the membrane. Nanoclustering of the receptor on the membrane is observed, indicating successful transfection and correct incorporation of the transmembrane domain. (**E**) Representative image of a MerTK transfected HeLa cell, permeabilized and stained for MerTK intracellularly. No nuclear localization of MerTK is observed. (**F**) Representative image of permeabilized Hek293 cells stained for MerTK. Cells endogenously express MerTK, but it is not found in the nucleus. (**G**) Representative image of a Hek293 cell after transfection with MerTK to induce overexpression of the protein. Even though a clear increase in the intracellular MerTK level is obtained, nuclear localization is not observed.

**Figure S4:**
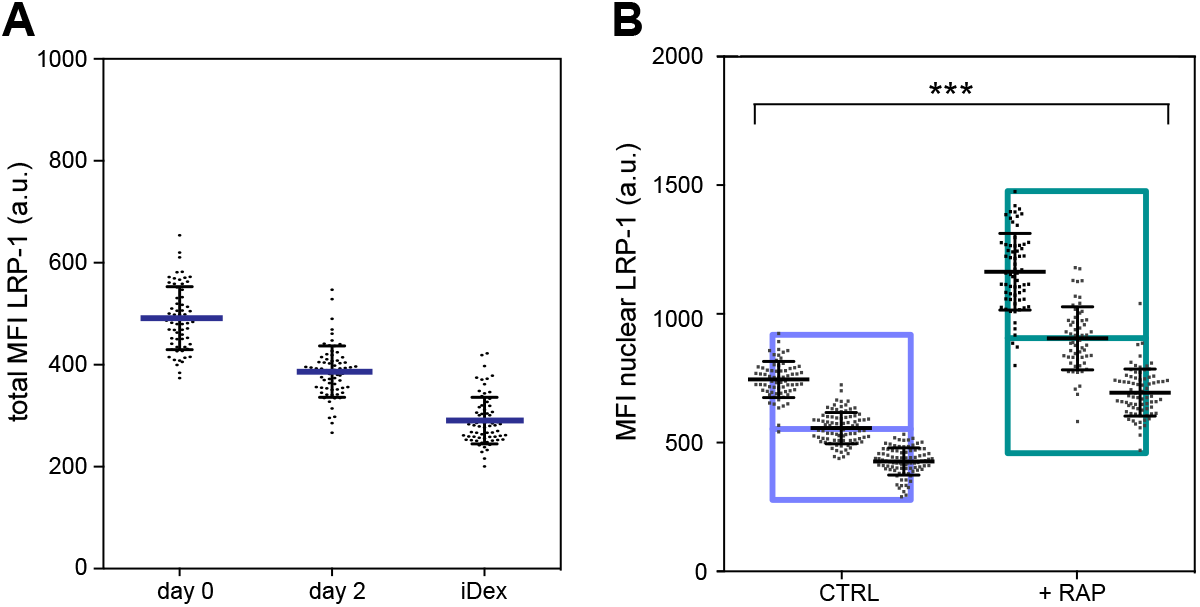
Total LRP-1 expression levels. (**A**) MFI of LRP-1 in the entire cell (membrane - which did not exceed isotype control levels, cytosol and nucleus) over time in culture, at days 0, day 2 and iDexs (= day 4). Data from 3 donors. (**B**) MFI of nuclear LRP-1 with and without the addition of RAP, a ligand of LRP-1, to the culture. Monocytes were isolated normally, and RAP was added to one of the cultures after a few hours up to day 0, when the cells were harvested for imaging. Each spot represents a single nucleus; small plots in larger bars correspond to data from 3 different donors. Statistics was performed using the average value of each donor.

**Figure S5:**
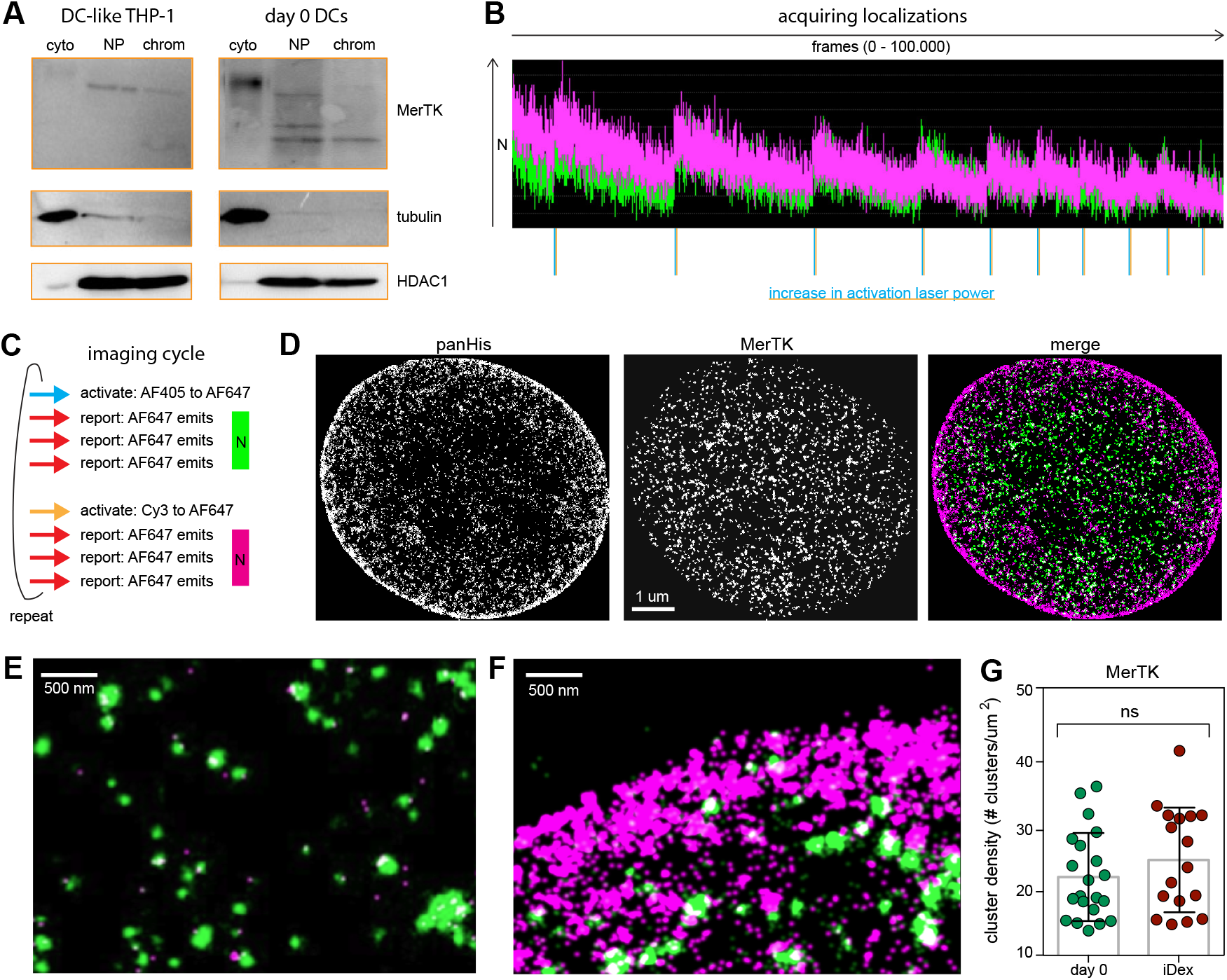
Super resolution STORM imaging of MerTK and chromatin. (**A**) Representative Western blot showing the relative abundance of MerTK in different cellular fractions (cytoplasm, nucleoplasm, chromatin-bound) in both DC-like THP-1 and day 0 DCs. Tubulin and HDAC1 staining indicate correct fractionation of the cells. (**B**) Representative imaging trace during STORM acquisition. The time expressed in number of frames acquired is plotted against the number of localization identified per frame (N). The vertical lines under the plot indicate when the activation laser power was increased, which is clearly seen in the peak increase in localizations registered. Over time the number of localizations is progressively reduced, indicating exhaustive imaging of all fluorophores. The traces also show that the number of localizations for both colours (MerTK in green and panHis in magenta) are comparable, validating the use of 2-color STORM imaging with minimal risk for bleedthrough. (**C**) Schematic representation of one imaging cycle: First, AF405 staining MerTK is activated by a pulse of the 405 nm laser line, and photons are transferred to the attached reporter dye AF647. Then, 3 pulses of the 647 nm laser line promote the emission of photons from AF647 and places the dye back into the dark state, until after the last pulse no localizations are recorded anymore. This makes the way free to start imaging the next colour without crosstalk. Cy3 staining panHis is activated by a pulse of the 561 nm laser line, and photons are transferred to attached reporter dye AF647. Then, 3 pulses of the 647 nm laser line force the emission of photons from AF647, and places the dye back into the dark state, ready for the next imaging cycle. (**D**) Pixelated binary reconstructed STORM images of both panHis and MerTK, and the merge of both channels (green for MerTK and magenta for panHis). Pixel size corresponds to the position accuracy, namely 20 nm. These binary images are used to calculate the colocalization between both colours. Binary pixelated images are required for this correlation analysis since raw localizations with an exact *x,y* positions will never perfectly colocalize and are therefore are not suitable for determination of the degree of colocalization between MerTK and panHis on a pixel to pixel basis. (**E**) Representative reconstructed STORM image of a cytosolic area in which no colocalization between MerTK and panHis is expected, as panHis labels the nucleus. MerTK vesicles show signal almost exclusively resulting from the AF405/AF647 dye pair (with only 2% of crosstalk with the panHis signal). (**F**) Representative reconstructed STORM zoom-in image on the nuclear periphery where the signal arises almost exclusively from we find the Cy3/AF647 dye pair labelling panHis. (**G**) Quantification of the number of MerTK nanoclusters per μm^2^ in the nucleus of both day 0 DCs and iDex DCs. No significant difference is observed. The data was obtained from the MerTK channel of the dual colour STORM images. Each dot corresponds to an individual nucleus. Data from 3 different donors. The average value per cell is depicted.

**Figure S6:**
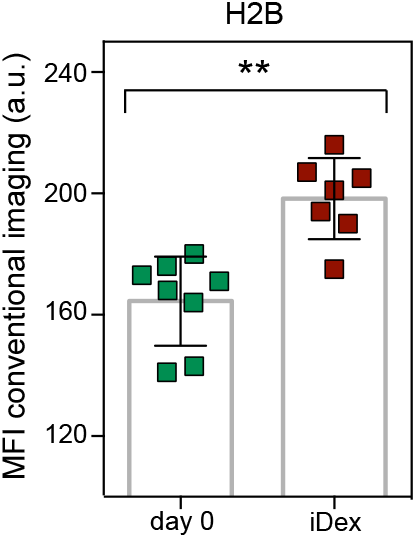
Expression levels of H2B in the nucleus of day 0 vs iDex DCs. (**A**) MFI of nuclear H2B in day 0 vs iDex DCs. Conventional images were taken before STORM acquisition of the same cells. This shows, together with Figure 5E,F, that the global increase in localizations per nanocluster and nanocluster density is indeed also reflected by a total increase of expression levels of H2B in the nucleus.

**Figure S7:**
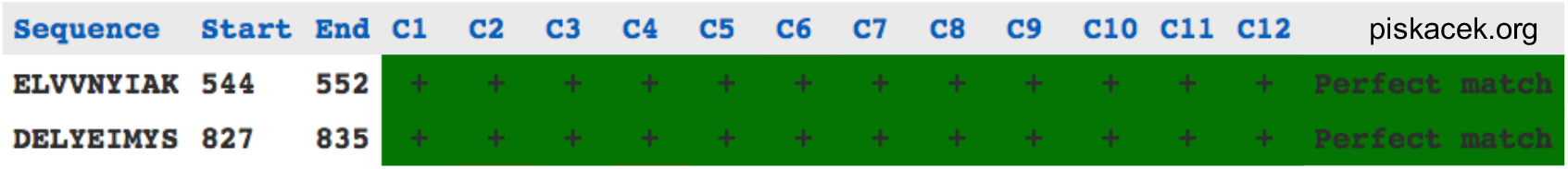
Prediction of 9aaTAD domains in MerTK sequence. Prediction of putative 9aaTAD domains in the aminoacid sequence of MerTK. Prediction was performed using an algorithm developed by Piskacek et al (Piskacek et al., 2016, 2007). The algorithm was accessed through their website Piskacek.org and the settings recommended for mammalian cells were applied (moderately stringent pattern).

